# Shaping hydrogel bioinks into 3D, multiscale, perfusable models using multimodal printing

**DOI:** 10.64898/2026.01.29.702588

**Authors:** Puskal Kunwar, Arun Poudel, Ujjwal Aryal, Zachary J Geffert, Daniel Fougnier, Ameya Narkar, Kairui Zhang, Alex Filip, Pranav Soman

## Abstract

Despite technological advances, the fabrication of multiscale, multi-material, and topologically complex 3D structures using soft hydrogel bioinks remains a challenge due to the inherent trade-offs between print size/resolution, bioink properties, and design complexity. In this work, we combine additive (macroscale) digital light projection (DLP) mode with subtractive (microscale) two-photon ablation (TPA) mode with multi-material exchange capability. We identify ideal hydrogel bioink formulations that are compatible with both DLP and TPA modes of processing. Technical challenges related to multimodal fabrication such as alignment of multiscale topologies to facilitate seamless media perfusion, soft-hard multi-material printing to facilitate handling of mechanically weak hydrogel constructs, and hydrogel swelling during printing, were resolved. To highlight the novelty of this hybrid platform, we fabricated centimeter-scale bioink constructs with embedded microscale perfusable topologies that cannot be achieved by isolated use of either DLP or TPA modes. This includes simpler microfluidic chips with independently perfusable microchannels to more complex 3D constructs with embedded, multiscale, perfusable dual-fluidic circuits that mimic the alveoli-capillary interface, or microfluidic chips with endothelialized microchannels. The unique ability of this multimodal platform to mimic *in vivo*-like multiscale complexities can be potentially used to develop next-generation organ-on-chips.

## Introduction

As compared to Nature’s ability to generate elegant multiscale 3D perfusable topologies using soft materials, current fabrication methods remain quite limited. Hydrogels, water-swollen polymer networks that mimic many aspects of native tissues, have been widely used in biomedical engineering, however shaping them into multiscale 3D constructs with high design complexity remains a challenge. Since structural complexities of native tissue are inextricably linked to their biological function, having the ability to fabricate anatomically accurate models using hydrogel bioinks will lead to new organ-on-a-chips for studying developmental biology, drug testing, disease modeling, and personalized medicine. Current methods of shaping hydrogels, based on extrusion, inkjet, and light-crosslinking, are limited in terms of their scale-range, design complexity, and choice of bioink formulations.^1-6^ For instance, complex hollow topologies with extrusion printing requires the use of sacrificial material or a support bath, however the resolution is typically lesser than 100 µm.^7-24^ On the other hand, Digital Light Projection (DLP) and their advanced variants can print complex 3D designs at higher speeds, however their resolution is typically ≥ 50µm.^3, 25-32^ For printing hollow topologies below 50 µm, two-photon ablation (TPA) is the primary option,^33-34^ however low print speeds make this unsuitable for making multiscale constructs. New multimodal printing platforms have been developed to mimic multiscale biological complexity, however this field remains in its infancy.

Due to the technical challenges in integrating specific hardware/software associated with different methods into one machine, most multimodal methods rely on sequential, unidirectional processing steps to generate composite constructs using separate machines or processes. For instance, dipping 3D printed structures into cell-laden solution, volumetric additive manufacturing (VAM)-enabled overprinting of structures around pre-fabricated electrowritten scaffolds, melt electrowriting followed by inkjet bioprinting, extrusion bioprinting followed by optical-patterning via DLP, vat photopolymerization combined with field-based patterning are few examples to generate multi-scale composites.^35-40^,^41-43^ A sequential process of crosslinking either solid or hollow microstructures within pre-fabricated microfluidic chips using two-photon polymerization (TPP) has also been developed, although removal of such structures from the chips or scaling-up beyond the dimensions of the chip is not possible.^44-45^ Subtractive methods such as 2PA have also been combined with additive methods such as VAM to fabricate multi-scale constructs,^46^ however strict limits on working depth (≤500 μm) necessary for TPA limits its utility. Thus, due to often contrasting requirements related to material processing and feature sizes along with lack of integrated control systems to precise align multimodal features, there are limited examples of hybrid machines that can bidirectionally switch between printing/processing modes to develop multiscale and/or multi-material constructs. For instance, Hybprinter combines molten material extrusion (∼350 μm) with DLP-SLA (∼35 μm),^47^ while Dual 3D bioprinter that can be switched between extrusion-printing and SLA-crosslinking modes to print PLA with cell-laden gelatin methacrylate (GelMA).^48^ Large area projection micro-stereolithography (LAPμSL) combines galvanometric mirror with a scanning lens to generate centimeter scale metallic structures with microscale resolution.^49^ Hybrid Laser Platform (HLP)^50-51^ integrates three modes of operation into one machine. HLP can switch between DLP, subtractive TPA, and additive TPP modes to generate centimeter-sized constructs with a resolution of ∼1-3 µm. Another example is Multipath projection stereolithography (MPS)^52^, which can switch between high- and low-resolution optical paths and can print centimeter-sized chips (3 cm × 6 cm) with a resolution of ∼10 μm. Despite these advances, LAPμSL, HLP, and MPS have never been utilized with difficult-to-print hydrogel bioinks. Thus, present hybrid 3D printing remains in its infancy, especially in the context of difficult-to-process bioinks, where contrasting requirements of printing fidelity (resolution, 3D design complexity) and biocompatibility must be met.

With the long-term goal of making next-generation biomimetic tissue models, we report the use of DLP-TPA multimodal printing using hydrogel-based bioinks. After optimizing bioink formulations that are compatible with both DLP and TPA modes, key technical challenges related to alignment of multiscale hollow features, swelling during printing, handling soft bioink constructs, were addressed. Finally, we designed and fabricated 3D centimeter-sized structures with microscale, topologically perfusable microchannels with increased complexities.

## Results

### Screening cell-adhesive hydrogel bioinks that are compatible with DLP and TPA modes

Before shaping bioinks into complex multiscale topologies, we screened 13 hydrogel formulations to identify ideal bioinks that can be additively printed using DLP mode, ablated using TPA mode, exhibit high cell adhesion, and sufficient robustness for handling. (**SI-1**). All bioink formulations were prepared using 1wt% LAP (photoinitiator) and 0.01 wt% Tartrazine (UV absorber). We found that widely used Gelatin methacrylate (GelMA) was not compatible with the TPA mode. For instance, fs-laser scanning (800nm, 1.2W, 100µm/s) within DLP-crosslinked GelMA (10%, G’100#1) resulted in a series of micro-bubble explosions with uneven removal of materials. **(SI-2A)** Then we tested 6 kDa poly(ethylene glycol) diacrylate (PEGDA), a biocompatible, low-viscosity, widely used material for DLP-printed hydrogels. Our results show that well-defined and unobstructed microchannels can be generated deep (∼1.5mm) within DLP-crosslinked PEGDA 6k gels due to minimal light scattering **(SI-2B)**. Since PEGDA 6k is inherently bioinert, we tested new formulations by adding integrin-binding motif RGD or adhesive proteins. Results show that by addition of methacrylated collagen remains compatibility with DLP-TPA modes, but cell adhesion remains poor **(SI-2C)**. Adding methacrylated RGD to PEGDA 6k showed uniform cell spreading by day 7, however high swelling (5x) resulted in detachment of cells by day 10. **(SI-2D)** To reduce swelling, crosslinkers MBAA and PEGDA 700 were added. Results showed that formulation with 2.4% MBAA **(#8, SI-1)** show high cell adhesion with confluent monolayer formation by day 3, although high-cost of RGD limits its scalability. **(SI-3A)** As an inexpensive replacement for RGD, we also tested 5 formulations (**#9-13, SI-1**) by adding varying amounts of GelMA (that possess cell-binding motifs to support cell adhesion) to the base formulation composed of PEGDA 6k. We also added PEGDA 700 to improve print fidelity and reduce swelling. Results show that these formulations are compatible with both DLP and TPA modes and show robust attachment and confluence of endothelial cells 48 hours after seeding. **(SI-3B)** Based on these screening results, bioink formulation **(#12: G50P40PL10, SI-1)** was selected for further studies, and in some cases, results were compared with inert PEGDA 6k bioink **(#2: P’100, SI-1**). This ideal bioink formulation can be additively printed using DLP mode, reproducibly ablated via TPA mode, and support high cell adhesion.

### Macro-to-meso scale printing of cell-adhesive bioink using additive DLP-mode

Figure **SI-4** shows the design and operation of additive DLP mode and subtractive TPA modes.^50^ The additive DLP mode consisted of a CW laser beam spatially modulated via a Digital Micromirror Device (DMD) to project a continuous sequence of light patterned through a transparent poly(dimethylsiloxane) (PDMS) bottom window. **(Figure 1A)** The use of an oxgen-permeable PDMS film creates an inhibition zone, also known as a dead zone, rich in radical-scavenging oxygen, which yields a thin uncured bioink layer between the PDMS film and the crosslinked hydrogel layers. During printing, the crosslinked part moves upwards, creating suction forces that renew the uncrosslinked bioink in the fabrication window. Further, the dead zone prohibits the adhesion of the crosslinked part to the substrate of the bioink bath, thereby preventing adhesion-related defects in this technique of 3D printing. Herein, we studied the deadzone to determine its thickness using cell-adhesive (**#12**) and cell-inert (**#2**) bioinks (**Figure 1B**). For the laser dosage used in this work, the dead-zone thickness ranges from 35 μm to 95 μm, exhibiting an increasing trend as the laser dosage decreases. This shows that additive DLP mode can be used to print structures in a continuous fashion using both inks. Next, we studied the lateral and axial resolution of the printed structure. Briefly, the lateral resolution was determined by a grid pattern with lines of 1-pixel to 12-pixels with an exposure time of 2 sec and laser intensity of 280 mW in the fabrication area **Figure 1C (SI-5)** displays a grid pattern printed on a glass substrate, demonstrating a resolution of 70 µm for 6 pixels, along with the field of view of 1 cm^2^. During the fabrication development process, grid patterns printed with 1-pixel, 2-pixel, 3-pixel, and 5-pixel lines were washed away. However, the 6-pixel line achieved a line width of 70 µm at a laser power of 280 mW. At 225 mW laser power, even 12-pixel lines resulted in under-crosslinked structures. This was achieved using a laser power of 280 mW and an exposure time of 2 seconds **(Figure 1D)**. Further printing of circular channels demonstrated the successful fabrication of 200 µm channel sizes, enabling the creation of complex structures, channels, overhangs, and undercuts **(Figure 1E)**. Two complex structures were then printed using cell-adhesive bioink **(#12, SI-1)**. The vase structure shown in **Figure 1G** was printed at the speed of 36.8 mm/hr with a slice thickness of 50 µm and this took 13 minutes to print 0.8 cm structure. Using the same printing conditions, another 3D structure resembling the alveoli-capillary interface found in human lungs was also printed **(Figure 1H)**. This structure, with a minimum feature size of ∼50 µm, has two sets of independent fluidic circuits perfused with two different flood color dyes. In this work, tartrazine dye was employed as a photo absorber to reduce curing depth, enhancing the z-resolution to 165 µm. The dye can be easily washed out of printed hydrogel construct by simply dipping the printed structure in water **Figure 1H)**.

**Figure 1.**
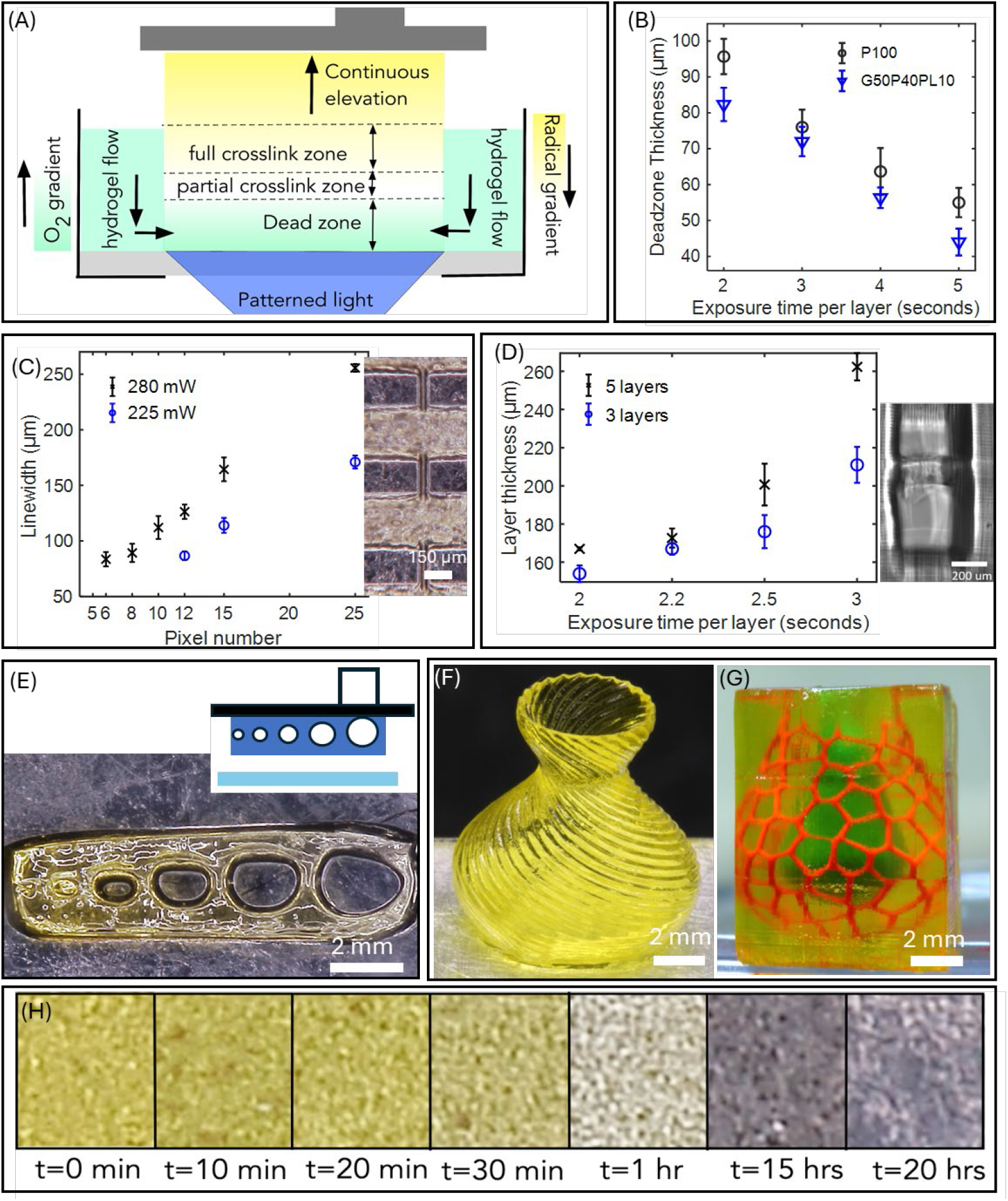
(A) Schematic of DLP-mode process. (B) Evaluation of dead zone thickness using bioink formulations #12 (G50P40PL10) and #2 (P100). (C) Plots showing a lateral (XY) resolution of 70 μm (D) Plot showing a vertical (axial, Z) resolution of 165 μm (E) (i) Schematic and associated printed structure showing an array of horizontal channels (F) Vase structure printed using bioink formulation #12 at a speed of 36.8mm/hr with a layer thickness of 50 μm. (G) Alveoli-like structure printed using bioink formulation #12 showing two distinct microfluidic circuits with different dye colors. (H) Snapshots of images showing the removal of tartrazine from printed structure after incubating them in DI water.

### Microscale ablation of crosslinked cell-adhesive bioink constructs using TPA mode

The subtractive TPA mode consists of a spatially cleaned, collimated, expanded fs-laser directed via an objective lens to enable ablation of hollow features within the previously crosslinked layers. **(SI-4B)** In TPA, the fs-laser beam is spatially cleaned and expanded by using 4x lens system and focused through an objective lens to ablate voids inside previously cross-linked layers with minimal collateral damage to the surrounding region. With hydrogel bioinks, several coupled mechanisms such as two-photon absorption, plasma generation, and subsequent cavitation bubble formation are observed (**Figure 2A)**. When fs-laser irradiation is applied to pre-crosslinked bioink slabs, varying the laser parameters enables access to different processing regimes. At lower intensities, localized densification occurs, producing a measurable change in refractive index without physical material removal while at higher intensities, plasma formation triggers pressure waves that drive cavitation, leading to bubble formation and expansion within the hydrated bioink slab to a certain size based on the laser fluence. These bubble collapse when the laser beam is turned off as the bioink matrix relaxes after the bubble collapses resulting in a decrease in average channel diameter **(Figure 2B, Video 1)**. With bioink formulation #12, we found that the maximum diameter of the bubble for the 280 mW laser power was observed to be 30 µm. When the laser is turned off this bubble collapse to 25 µm. The focal spot of the laser beam, determined using the Rayleigh criterion and focused through an objective lens with a numerical aperture of 1.2, is approximately 0.27 µm. These bubble features can expand ∼100 times larger than the estimated laser spot size within the crosslinked bioink. During scanning it is observed that the bubble moves with the laser focal point creating a void channel and surrounded by a possibly over-crosslinked shell outline (**Figure 2C (i-iii)**), likely induced by the fs laser exposure during ablation. **(Video 2)**. We utilized this technique to generate a complex 3D structure test pattern (human alveolus alike structure) within previously crosslinked slab structure printed using bioink formulation #12 **(Figure 2C (iv)**). The 3D hollow structure, mimicking the capillary network that surrounds alveoli, was generated ∼100μm below the surface and extended 200 μm further into the gel. This structure was imaged using a bright field microscope. This result shows that the TPA mode can fabricate complex, perfusable 3D microfluidic network that mimics in vivo structural complexity.

**Figure 2.**
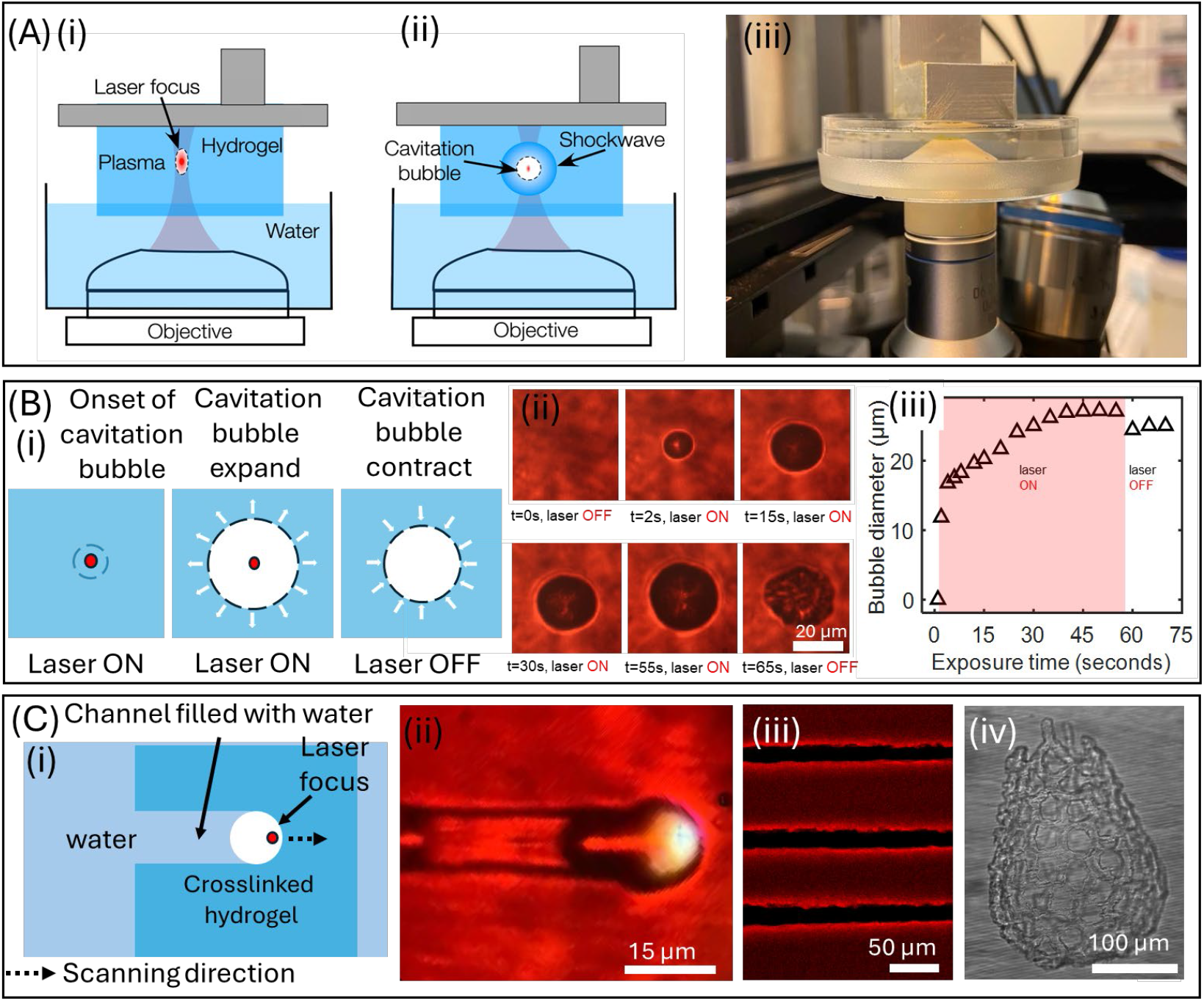
(Ai,ii) Schematic of TPA in previously crosslinked bioink structure. (iii) Picture of the experimental setup used for TPA showing a drilled Petri dish affixed to the immersion objective lens. (B) (i) Schematics (ii) experimental results and (iii) plot showing the pressure cavitation dynamics during TPA within structure crosslinked using bioink formulation #12. (i) Schematic cartoon, (ii) Experimental results showing channel fabrication via cavitation in TPA mode. (iii) simple microchannels and (iv) 3D microchannel network mimicking capillary network around alveoli sacs in lung tissues.

### Fabrication of multiscale bioink constructs and multi-material composites using DLP-TPA

Bioink formulation #12 was then used to print 3D constructs with embedded microscale perfusable topologies using sequential and/or bidirectional use of additive DLP mode (resolution range ∼1 cm to ∼100 µm) and subtractive TPA mode (resolution range ∼50 µm to ∼3 µm). A typical workflow involves crosslinking of bioink layers using DLP mode followed by targeted fs-laser ablation within the crosslinked layer. This process can be repeated, and modes can be bidirectionally switched to realize the fabrication of hollow micro-features at virtually any depth. A washing step is necessary between mode switching to wash away any un-crosslinked resin. For instance, switching from DLP to TPA mode, fs-laser could initiate unwanted crosslinking of bioinks which can block the embedded perfusable topologies and modify solid features generated during previous steps. In this setup, resin vat is replaced with another vat filled with DI water; here printed structures are immersed in DI water and syringe was used to wash the embedded topologies to remove any uncrosslinked resin.

The first design consists of a microfluidic chip (total thickness = 1.5 mm) with two independent mesochannels (width=500 µm), each with inlet and outlet open wells, connected with an array of microchannels (lumen size of 10 µm) embedded 250 µm inside bioink structure. (**Figure 3A**). DLP mode was used to print the chip with embedded mesochannels while TPA mode was used to ablate the microchannels. Perfusability of embedded microchannels was verified by adding a solution containing red microbeads (FluoSpheres™, size-1μm) into one of the wells. Results show flow of microbeads, driven by both static hydrostatic pressure and capillary action, within the microchannels **(Video 3)**. The second design consists of 4 wells printed using DLP mode and two embedded microchannels separated by ∼10µm. (**Figure 3B**). Wells 1 and 3 are filled with dyes with yellow microspheres in left microchannel and green FITC-dextran in the right microchannel. The overall size of the chip is 6 mm x 6 mm part size and total printing time for these chips is less than 10 minutes.

**Figure 3.**
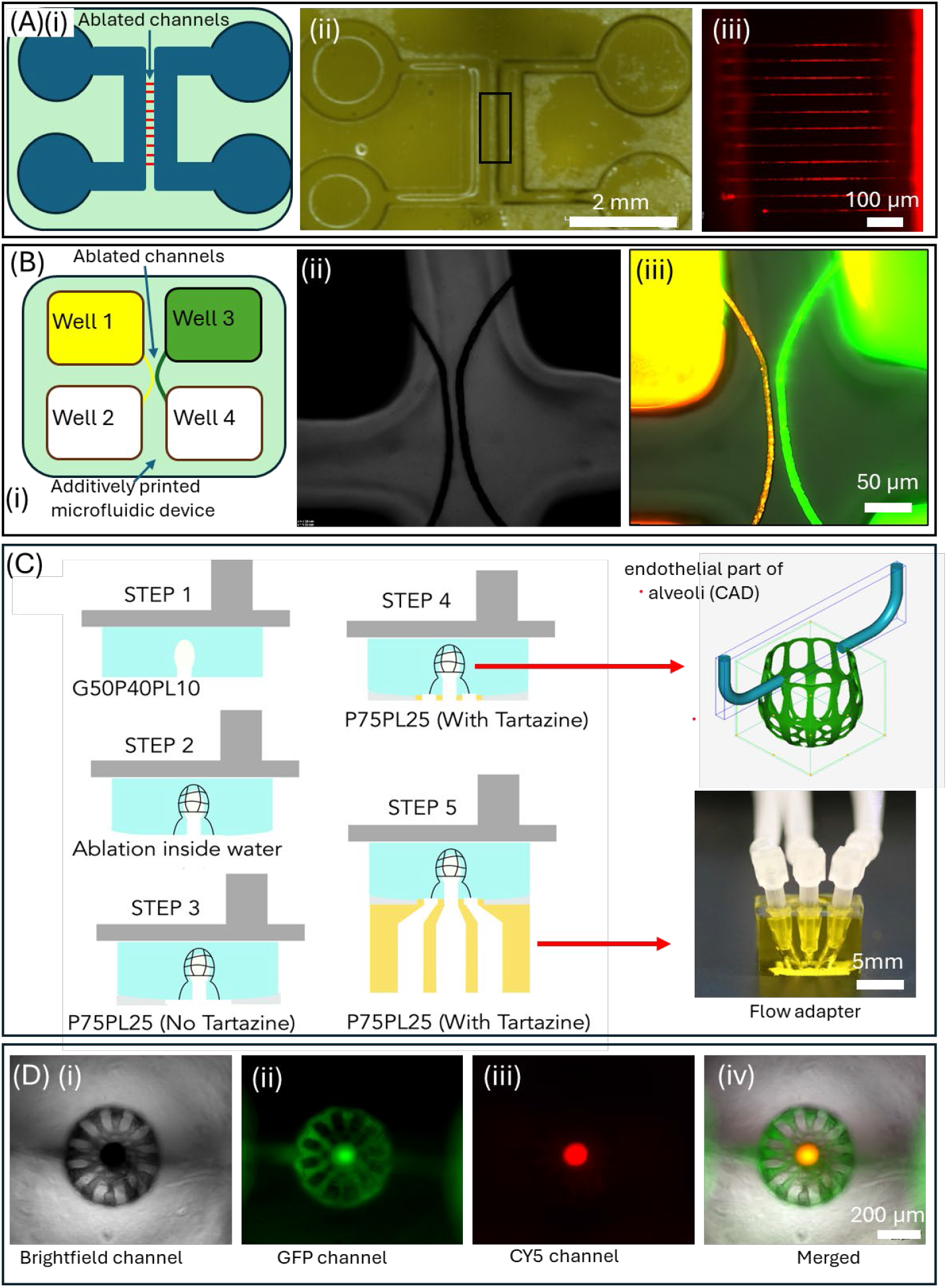
(A) (i) Schematic of the microfluidic chip using DLP-TPA modes. (ii) Fabricated microfluidic chips with meso-structures printed via DLP mode and microchannels via TPA mode. (iii) Fluorescence image showing flow of red fluorescent dye through ablated microchannels. (B) (i) Schematic of four well microfluidic chips with two independent channels. (ii,iii) Brightfield and fluorescence images showing perfusion of red and green dye in adjacent channels separated by a 10 μm gap. (C) Schematic of process flow showing bidirectional, multimodal, and multi-material printing to print alveoli-capillary structure. (D (i-iv)) Representative images showing alveoli-like structure with red and green dyes in alveoli-sac and capillary structures.

Design 3 was inspired by the alveoli-capillary interface found in lungs. In vivo, individual spherical alveoli (approximately 100-300 μm) are surrounded by a dense network of capillaries (approximately 10-30 μm). The 3D design complexity and the overall size of this structure brought forth new challenges such as swelling or de-swelling during mode switching, associated feature misalignments, and practical issues such as perfusion within capillary-size channel networks. We observed that there is a discernible difference in the ablation z-range between the water dipping objective lens and the air objective lens. We made the deliberate choice to employ a water dipping objective lens due to its capacity for achieving a greater penetration depth and ablation z range. This choice bears the advantage of reducing the frequency of transitions between additive and subtractive modes, as well as the interruptions during washing steps. However, we observed a gradual change in the dimensions of additively crosslinked structures during slow TPA mode, especially longer for complex alveoli-capillary designs. To correct for swelling-induced misalignment, we investigated this in more detail. Based on a layer-by-layer setup used for additive DLP mode, bioink swelling is constrained along the z-direction while swelling is more evident in the lateral dimensions. Swelling induced shape changes **(Figure S7**) were incorporated into the CAD model for all 3D complex topologies, which typically take longer times during TPA mode. Design 3 consists of a design that mimics the alveoli-like architecture found in lung tissues. Here, a section of the air sac was additively printed using the additive DLP mode, then the mode was switched, and a capillary-like network was ablated using TPA mode. DLP and TPA modes were switched back and forth to print the entire structure. (**Figure 3A, step 1-2**). Connecting these mechanically soft constructs to external perfusion flow required us to print a robust flow-adapter that is directly attached to the soft constructs. As noted earlier, time-consuming TPA mode results in a dome-shaped topology at the printing (lower) interface, **(SI-6B)** which makes is challenging to directly print the flow adapter directly. To flatten the domed surface, an initial layer was crosslinked using the P75PL25 formulation (P =10% PEGDA 6k, 90% PBS; PL =20% PEGDA 700 + 80% DI water) without tartrazine, to ensure higher penetration depths and flattening out of the domed interface. **(step 3**) Bioink formulation was switched again to P75PL25 with tartrazine to print the remaining layers that make up the flow adapter (**step 4-5**), as shown in **Figure 3C**. This ensures that hollow features of the adapter just above the alveoli-like topology remain open. Finally, two distinct fluorescent dyes of different colors (FITC-dextran, green, and red FluoSpheres™; 1μm) were introduced into independent circuits of alveoli like structure using a pipette, demonstrating the functional perfusion capabilities of the printed structure (**Figure 3D i-iv**). This approach highlights the versatility of hybrid printing in creating complex, multiscale, and perfusable biological architectures. The overall size of the structure with adapter structure was 10 mm x 10 mm x 8 mm, whereas the size of the alveoli was ∼500 µm and the ablated microchannels ranged from 10-50 µm. Please note the lumen size possible with TPA is 10x smaller compared to those possible with additive DLP mode. **(Refer to Figure 1G)**

Design 4 further increased the size and complexity of the constructs. To replicate the multiscale complexity found in the bronchial tree, which consists of bronchi that branch into progressively smaller passageways before terminating in tiny air sacs called alveoli, we designed a CAD model of a simplified bronchiole with three alveoli structures (∼100-300μm) independently surrounded by capillary-sized microchannels (10-30μm) using bioink formulation #12. **(Figure 4A)**. CAD model designed in SolidWorks was divided into two parts, one for DLP and one for TPA mode. The model used for DLP mode was sliced to generate digital masks for the DMD using a custom-written MATLAB algorithm. A series of bi-directional switching between DLP and TPA modes was used to print the structure. A crosslinking laser wavelength of 405 nm and a crosslinking laser power of 280 mW (2.17 mW/cm^2^) and exposure time of 2 seconds were used to selectively crosslink the prepolymer solution in an additive continuous fashion on a surface-modified glass coverslip. The subtractive mode, with an ablation laser wavelength of 800 nm, ablation laser power of 1200 mW, and scanning speed of 100 µm/s, was used to ablate microchannels surrounding the alveoli structure. The overall dimension of the fabricated bronchiole-like structure was 1 cm x 1 cm x 0.75 cm, and the single alveoli structure fits within a cube-frame size of 600 × 600 × 600 μm^3^ and ablated channel diameters ranging from 10 to 30 μm (**Figure 4A ii-iv**). The total printing time for the structure included 2 hrs and 25 mins for steps in TPA mode, 16 mins for all steps in DLP mode, and a combined transition time of under 9 min between steps. To facilitate perfusion within capillary network structures, the design was modified to allow gradual changes in the cross-section between microscale and mesoscale hollow features **(SI-7)**

**Figure 4.**
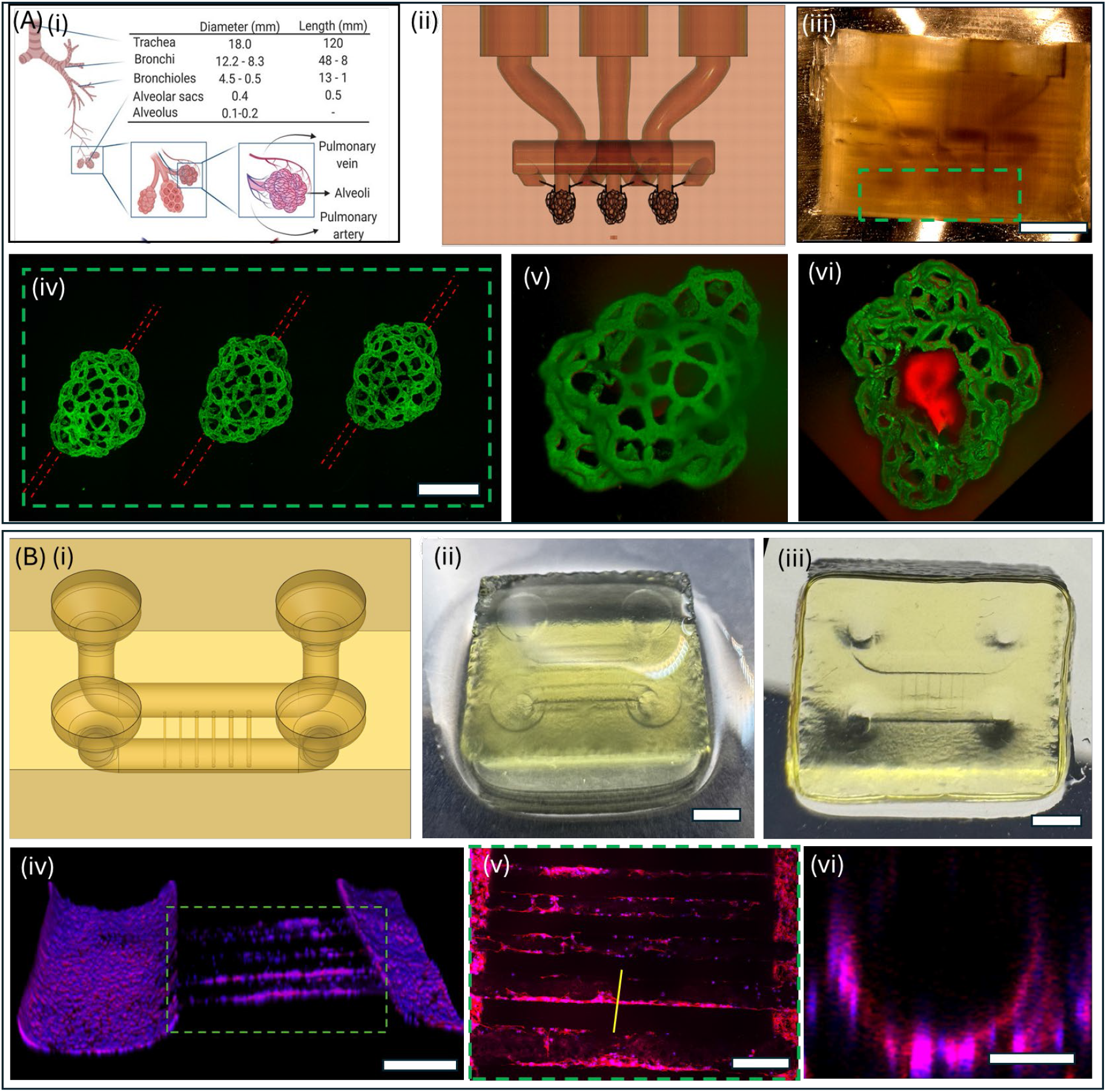
(A) (i) The hierarchical structure of the lung showing spherical alveoli sacs (∼200 μm) surrounded by capillary networks (∼10-30 μm). (ii) Simplified CAD file. (iii) Bright-field image of printed structure printed using DLP-TPA modes. (iv) fluorescent images of three capillary structure after perfusing FITC-dextran (Green) dye. (v,vi) top and bottom view of the alveoli sacs with red dye and capillary structures perfused with green dye. (B) (i) CAD of microfluidic chip with embedded microchannels (ii,iii) Bright-field top and bottom images of printed structure with an array of channels varying from 2mm to 10µm; Channels >100 µm printed using DLP mode while channels <100 µm are printed using TPA mode. (iv-vi) Confocal images of channels lined with 2H11 endothelial cells lining. (vi) cross section of the channel shown in (v).

Design 5 focused on generating multiscale vasculature-on-a-chip due to its potential impact for drug discovery, disease modeling, and personalized medicine. Bioink formulation #12 was used to fabricate a construct with an overall size of 1 cm x 1 cm x 0.5 cm, with a pair of inlet-outlet ports for easy pipetting of cell solutions. The pipetting well (2-mm diameter) mesoscale channels (1-mm in diameter), printed by DLP-mode, are connected by a series of microchannels (size range 10-50 µm) ablated using the HLP mode (**Figure 4 B (i-iii)**). For DLP mode, a laser intensity of 280 mW (2.17 mW/cm^2^), a layer thickness of 50 µm, and an exposure time of 2.5 seconds per layer, were used. For TPA mode, ablation of microchannels was performed using the laser intensity of 1200 mW and scanning speed of 100 µm/s. **Figure 4B (i)** displays a confocal image of the multiscale structure with a 2 H11 lining and zoomed section (**Figure 4B (ii)**), indicated by a green dotted rectangle, revealing channels with endothelial cell lining inside the ablated channels. The ablated channels were only partially covered with endothelial cells, likely due to the static media cell culture (**Figure 4B (iii))**. We anticipate that this condition can improve with the induction of media flow by incorporating a suitable bioreactor.

## Discussion

Multiscale printing of 3D perfusable geometries using biomimetic hydrogels holds great potential for a range of applications, from microfluidic systems to organ-on-a-chip. However, the generation of multiscale freeform designs, especially using difficult-to-process soft hydrogel bioinks, is not possible with any single fabrication method. For instance, despite advances in both additive DLP and subtractive TPA techniques, each method has limitations in terms of their print size, resolution, design complexities and speed. Combining different, yet traditionally mutually exclusive, modes of processing bioinks into a single hybrid platform can be the solution. At present, most hybrid strategies rely on separate machines, requiring printed parts to be moved from one machine to another, and often a trial-and-error approach is then used to optimize print fidelity. Here, we integrate DLP and TPA modes into a single platform and use 5 different chip designs to showcase this platform’s novel capability. A key initial challenge was to screen a library of readily available bioinks and select an ideal formulation to balance often contrasting requirements of printing fidelity in DLP mode, reproducible ablation of microchannels, and sufficient cell adhesion. Multi-material DLP-TPA was used to print adapters to handle printed soft bioink structures and to facilitate perfusion flow within embedded microchannels. The integrated DLP-TPA exceeds the resolution and design flexibility associated with individual methods and provides a new method to make hierarchical structures using bioinks. Compared to extrusion-bioprinting methods that rely on the removal of sacrificial materials to generate microchannels, or traditional TPA with limited laser penetrating depths, combination of sequential layer-by-layer additive printing with fs-laser enabled cavitation can selectively generate hollow microfeatures at any user-defined depth, as demonstrated by this work. Please note that independent use of either DLP or TPA modes cannot be used to generate such structures. We envision that such chips could be used to mimic biological interfaces such as the alveolar-capillary found in lungs.

This work can be improved in many aspects in future studies. For instance, slow ablation speeds with TPA mode limits its practical utility to ≤ 1 cm^3^; this scale is sufficient for making advanced tissue chips but not sufficient for larger tissue engineering applications. Use of galvanomirror-assisted scanning, highly photosensitive bioinks, or multi-focus digital holography could be used to increase TPA speeds, however possible challenges with integrating these methods into a single machine remain unexplored. We also observed that certain bioinks exhibit defect-free ablation while some do not, and the underlying reason for this is not clear. Systematic studies need to be carried out to establish a relationship between bioink formulation and fs-laser cavitation. The control algorithm developed in this work can process user-defined CAD models and generate a series of digital masks for additive DLP mode and scanning paths for the TPA mode. However, current semi-automated steps to align features made by the two modes can be further improved. A closed-loop feedback control system can be added to automate bioink handling, washing steps, and corrective changes to CAD design, to reduce manual intervention and increase reproducibility. For chip design 5, we demonstrate endothelization of microchannels via cell seeding on acellular bioink chips as a proof-of-concept. Future work with complex topologies will be necessary to generate multi-cellular biology-mimicking interfaces using cell seeding strategies and improved designs of inlet-outlet ports to perfuse media under hydrostatic pressure or by using a flow pump. To demonstrate that this hybrid approach can be extended to cell-encapsulation, we bioprinted a slab structure with through microchannels (lumen size of 10µm) using 10T1/2-laden bioink (#12) and cultured under static culture. Although encapsulated cells are viable with uniform distribution around the ablated channels, perfusion within the ablated channels was not observed **(SI-8)**. Thus, current setup and possibly bioink formulations need to be modified to make multimodal printing compatible with cell-laden hydrogels. Other improvements could include the use of tissue-specific architectural blueprints, the use of iPSC derived cells, custom bioreactors with optimized media formulations, and functional characterization of endothelial and stromal cells. Overall, multimodal printing platform is an initial step towards to goal of making in vivo like organ-on-chips.

## Methods

### Setup for multi-modal printing

The additive DLP mode prints structures by projecting a continuous sequence of patterned light images by a digital micromirror device through a PDMS window, which is an oxygen-permeable, light-transparent window. The continuous wave (cw) laser beam produces 405 nm laser beam (Topica), which is spatially cleaned and expanded using 4x lens telescope. The beam is then spatially modulated using a DMD which creates light patterns based on user-defined images by toggling micromirrors between ON and OFF states. The beam is directed towards the sample bath that selectively crosslinks photosensitive prepolymer into 2D layers with a specific thickness. The laser beam modulated by the DMD can be used to crosslink photosensitive hydrogels in both continuous and layer-by-layer manner, with a feature size ranging from ∼30 μm to 1 cm. A custom written MATLAB code is used to slice a 3D model of the structure and to obtain a series of DMD mask. In subtractive mode (Figure **SI-4A**), the fs-laser beam (Coherent Chameleon-Ultra Ultrafast Ti:Sapphire, 800nm, 150 fs pulse width, 80 MHz) is spatially cleaned and expanded using a 4× lens system, then focused through an objective to ablate voids within previously cross-linked layers via two-photon absorption (TPA). This subtractive mode enables sub-micron feature removal, depending on the hydrogel’s absorption properties. Although the same fs-laser can also operate in additive two-photon polymerization (TPP) mode to fabricate 3D structures with microscale precision, that mode was not used in this study. The longer excitation wavelength increases penetration depth according to Rayleigh criteria, and the non-linear TPA process restricts energy deposition to the focal volume of the ultrashort pulses, enabling high resolution and precise 3D control (**SI4 B**,**C**). This mechanism allows localized ablation of microscale voids and channels within cross-linked structures while minimizing collateral damage.

### Synthesis of LAP

2.85 mL dimethyl phenylphosphonite and 3.00 mL 2,4,6-trimethylbenzoyl chloride were allowed to react under a nitrogen blanket in a foil-covered flask with magnetic stirring for 18 hours. The next day, 100 mL of 6.25% (m/v) lithium bromide in 2-butanone solution was added to the flask and the mixture was heated to 50°C. Once heated, the reaction was allowed to proceed for 10 minutes. The flask was then promptly removed from the heating bath and allowed to cool to room temperature. The resulting white precipitate was collected via vacuum filtration and washed with four 100 mL aliquots of 2-butanone. Isolated LAP product was subsequently dried *in vacuo* for 7 days to remove residual solvent.

### Synthesis of PEGDA 6k

18 g of 6 kDa poly(ethylene glycol) (PEG) was dissolved in 100 mL anhydrous dichloromethane (DCM) and the resulting solution was cooled in an ice bath. Once chilled, 1.69 mL triethylamine was added to the solution under a nitrogen blanket and the solution was allowed to stir for 15 minutes. 0.975 mL acryloyl chloride was diluted in anhydrous DCM to a final volume of 15 mL. The resulting diluted acryloyl chloride was added dropwise to the reaction flask via addition funnel and the mixture was allowed to stir under a nitrogen blanket at 0°C for 30 minutes. The flask was removed from the ice bath, covered in foil, and the reaction was allowed to proceed overnight with constant stirring. The reaction mixture was transferred to a separatory funnel and neutralized with 16 mL of a 2 M potassium carbonate solution with frequent venting to release CO_2_gas. Once gas evolution ceased, the resulting emulsion was allowed to separate on a ring stand for at least 4 hours. The bottom-most organic layer was collected and dried with anhydrous sodium sulfate. Excess sodium sulfate was removed via vacuum filtration. PEGDA was precipitated from the dried solution via dropwise addition into a 10x volume of hexane (approximately 1 L). The product was isolated via vacuum filtration and washed with 3 aliquots of hexane, followed by 3 aliquots of -80°C diethyl ether. Purified PEGDA product was dried *in vacuo* at room temperature for two days and stored at -20°C.

### Synthesis of GelMA

10 g of type A porcine gelatin was dissolved in 100 mL 1x phosphate buffered saline and heated to 50°C under a nitrogen blanket with constant stirring. Upon complete dissolution, 8 mL methacrylic anhydride was added via addition funnel at a rate of 0.5 mL/min. The reaction was allowed to proceed at 50°C under a nitrogen blanket with vigorous stirring for 3 hours. An additional 200 mL of PBS was added to the reaction mixture after the reaction mixture cooled to room temperature. The crude product was isolated via centrifugation 1000xg for 10 minutes, resulting in a viscous pellet. The product was collected and dialyzed in 12-14 MWCO dialysis membrane against 3.5 L of deionized water at 50°C. Dialysate solution was exchanged with fresh DI every 24 hours for 3 days. The pH of the product was adjusted to 7.0-7.4 using 0.2 M sodium hydroxide prior to freezing at -80°C overnight. Frozen samples were lyophilized to complete dryness before storage at -20°C.

### Rheological characterization

TA Instruments Discovery Hybrid Rheometer (DHR-3) with a temperature-controlled lower Peltier plate (TA Instruments, New Castle, DE, USA) was used for rheological measurements. Storage and loss moduli of photo-crosslinked hydrogel samples were measured via frequency sweep from 0.1 to 100 Hz at 25°C using 8 mm parallel plate geometry. Cylindrical photo-crosslinked hydrogel samples were prepared via exposure to 405 nm light patterned using an 8 mm diameter circular mask in a 1 mm deep PDMS well. Resulting hydrogel samples were 8 mm in diameter and 1 mm thick. All rheological measurements were performed in triplicate unless otherwise noted.

### Cell adhesion screening, and cell seeding within chips

The 2H-11 endothelial cell line, originally isolated from the axillary lymph node of an adult male mouse, and 10T1/2 fibroblast cell line, isolated from mouse embryo, were bought from American Type Culture Collection (ATCC, Manassas, VA). 2H11 cells were cultured in a cell growth media consisting of a Dulbecco’s Modified Eagle Medium (DMEM, GIBCO#11965-092) supplemented with 4 mM L-glutamine, 4500 mg L?^1^ glucose, 1 mM sodium pyruvate, 10% heat-inactivated fetal calf serum, and 1% penicillin–streptomycin. 10T1/2 fibroblast cell**s** were cultured in a cell growth media consisting of a Basal Medium Eagle (BME, GIBCO#21010-046) supplemented with 10% FBS, 1% penicillin/streptomycin and 1% GlutaMAX (GIBCO#35050-061). All the cells were cultured in T75 vented cell culture flasks and were maintained at 37 °C in a humidified incubator with 5% CO2. For cell seeding within the chip design 5, the luminal surface were surface functionalized using collagen. Rat tail collagen (Ibidi, Germany) was diluted using a 20 mM acetic acid solution to achieve a concentration of 0.5 ml/ml, and this solution was pipetted into the wells. After 30min of incubation, collagen solution was aspirated from the wells and the channels, followed by a triple rinse with PBS. Then, 2H11 cell solution (10 M/ml) was introduced into all four wells for 15 min to facilitate cell adhesion. Then, the cell solution was removed, fresh media added into the chip, and the chip was cultured in a static culture condition (37°C, 5% CO_2_ and 90% humidity) with media changed every 2 days.

### Imaging

To visualize cellular morphology, the cells were fluorescently stained for f-actin and nuclei and imaged using an LSM 980 confocal microscope (Zeiss, Germany). The same microscope was employed to capture images of the alveolar structures formed within the hydrogel. Brightfield and fluorescence images of some cellular and non-cellular constructs were obtained using an inverted microscope (Zeiss, Germany) and HIROX microscope. Line profiles of the structures were plotted and full width at half maxima (FWHM) was recorded to obtain the linewidth. The linewidth of the structures was averaged from 5 line profiles. To assess perfusion, three fluorescence solutions were used: FluoSpheres™ Polystyrene Microspheres (1 μm) yellow-green (505/515) and red fluorescent (580/605), and FITC-dextran (40kDa MW), all purchased from Invitrogen.

### TPA of microchannels within DLP-printed cell-laden constructs

DLP mode was used to crosslink a solution of C3H10T1/2-laden bioink (#12, 130 µl, 1 × 106 cells/ml) into a rectangular slab geometry (1mm high, 20 layers, layer thickness = 50 µm, 2.17 mW/cm^2^; 3s/layer). TPA was performed ∼100µm inside the crosslinked slab using a laser power of 1200mW, a scanning speed 100 µm/s at 800 nm wavelength. Then, constructs were rinsed with PBS for 5 minutes and cultured under standard laboratory conditions.

## Supporting Information

Additional figures and figure captions (SI-1 to SI-7), and caption for three Video files (V1 – V3).

## Acknowledgement

Financial support for this project was provided by the National Institutes of Health (R21DK136083, R21EY036189, R01AR083466), the Syracuse University Collaboration of Unprecedented Success and Excellence (CUSE) program.

