## Supplementary material for "Shaping hydrogel bioinks into 3D, multiscale, perfusable models using multimodal printing": SI Information

This file contains the following:

- Supplementary figures and figure captions (SI-1 to SI-8).
- Caption for Video files (V1 – V3)

| # | Bioink formulation | Print fidelity with additive DLP-mode | Ablation fidelity with subtractive TPA-mode | Biocompatibility or Cell adhesion |
|---|---|---|---|---|
| 1 | G'100 | ++ | + | +++++ |
| 2 | P'100 | +++ | +++++ | NA |
| 3 | P'90MC10 | ++++ | +++++ | NA |
| 4 | P'80MC20 | +++ | +++++ | + |
| 5 | P'70MC30 | + | NA | NA |
| 6 | P'80PL20R | +++++ | ++++ | ++ |
| 7 | P'80PL20RM1.2 | +++++ | +++ | ++++ |
| 8 | P'80PL20RM2.4 | +++++ | +++ | ++++ |
| 9 | G10P80PL10 | +++++ | +++++ | + |
| 10 | G20P70PL10 | +++++ | ++++ | ++ |
| 11 | G40P50PL10 | ++++ | ++++ | ++++ |
| **12** | **G50P40PL10** | **++++** | **++++** | **+++++** |
| 13 | G60P30PL10 | +++ | +++ | +++++ |

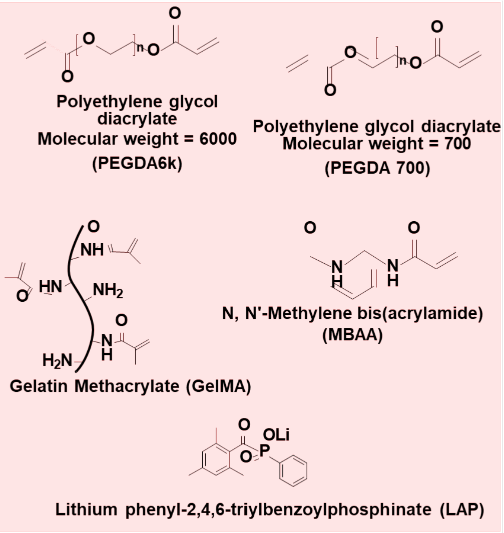


**SI-1.** Table showing the various bioink formulations tested, molecular structures of the base hydrogel, crosslinker, and photoinitiatorand their quanlitative performance in terms of being compatible with DLP-mode, TPA-mode, and cell-adhesion. All formulations were prepared in PBS containing 0.5% LAP as the photoinitiator and 0.01% Tartrazine as the photoabsorber. Bioink formulation #12 was chosen for most of this work. Qualitative screening of suitable bioink was performed using 2H11s, widely used immortalized murine vascular endothelial cell line.

**Nomenclature:** Letter (material) + number (% volume in the final formulation)

G' = 10% GelMA + 90% PBS; G =7.5% GelMA + 92.5% PBS

P' =15% PEGDA 6K + 85% PBS;

P =10% PEGDA 6K, 90% PBS;

PL =20% PEGDA 700 + 80% DI water;

MC = 20% Methacrylate collagen + 80% PBS;

R = methacylated RGD, M= MBAA (1.2% & 2.4%);

*All formulation were prepared in PBS with 1% LAP photo-initiator and 0.01% Tartarzine photo-

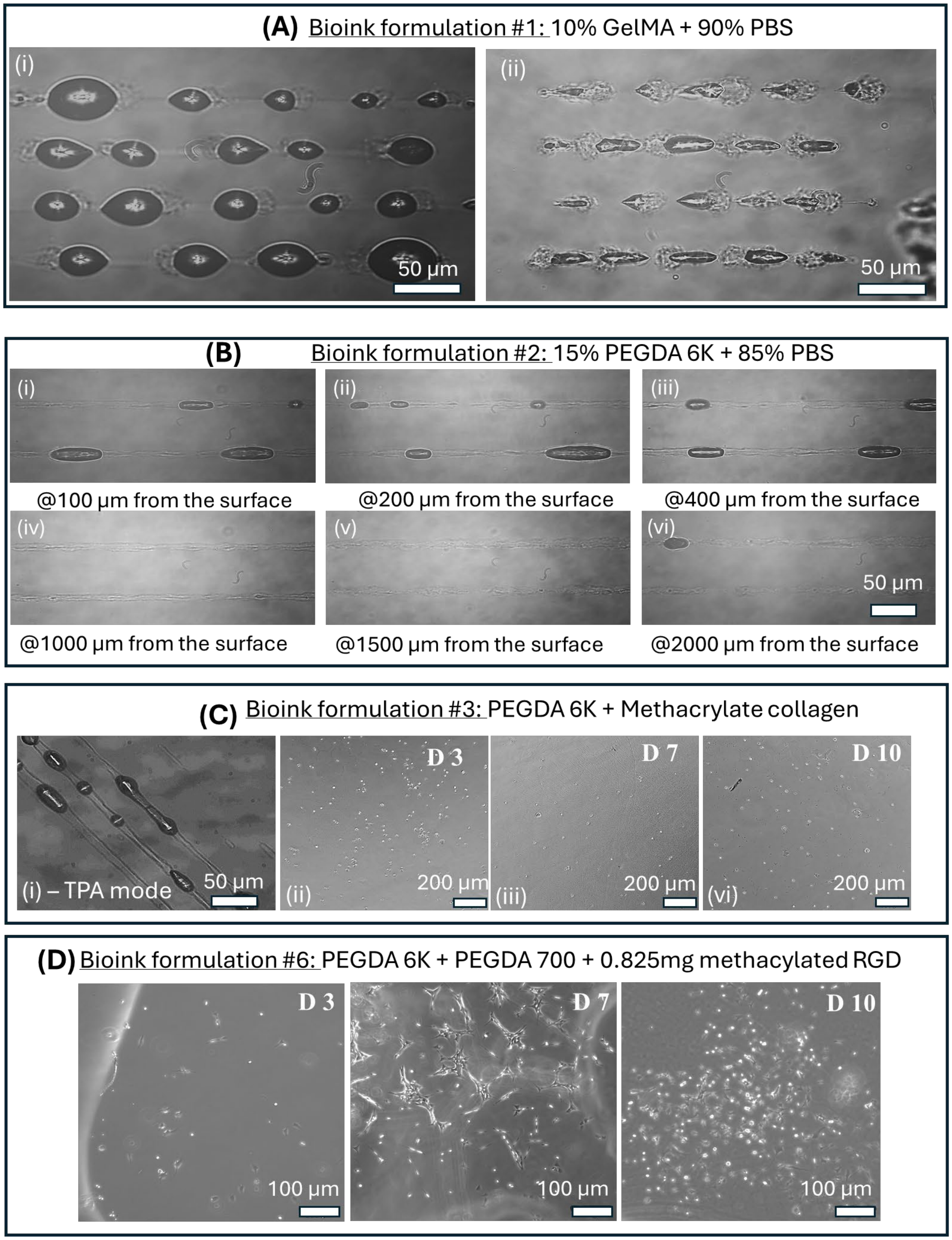


**SI-2.** (A) TPA-mode at depths of 100μm (i) and 200μm (ii) inside crosslinked GelMA slab (bioink formulation #1); here uncontrolled microexplosions prevent the formation of a fully perfusable channel (B) TPA-mode at varying depths inside crosslinked bioink slab (formulation #2); here fully perfusable channels are generated. (C) (i) TPA-mode in crosslinked slab using bioink formulation #3, (ii-iv) Cell adhesion on Days 3, 7 and 10. (D) Cell adhesion on Days 3, 7 and 10 using bioink formulation #6. **Note:** TPA mode was used with a power of 1.2W and scanning speed of 100 μm/s. Qualitative screening was performed using 2H11 cell line.

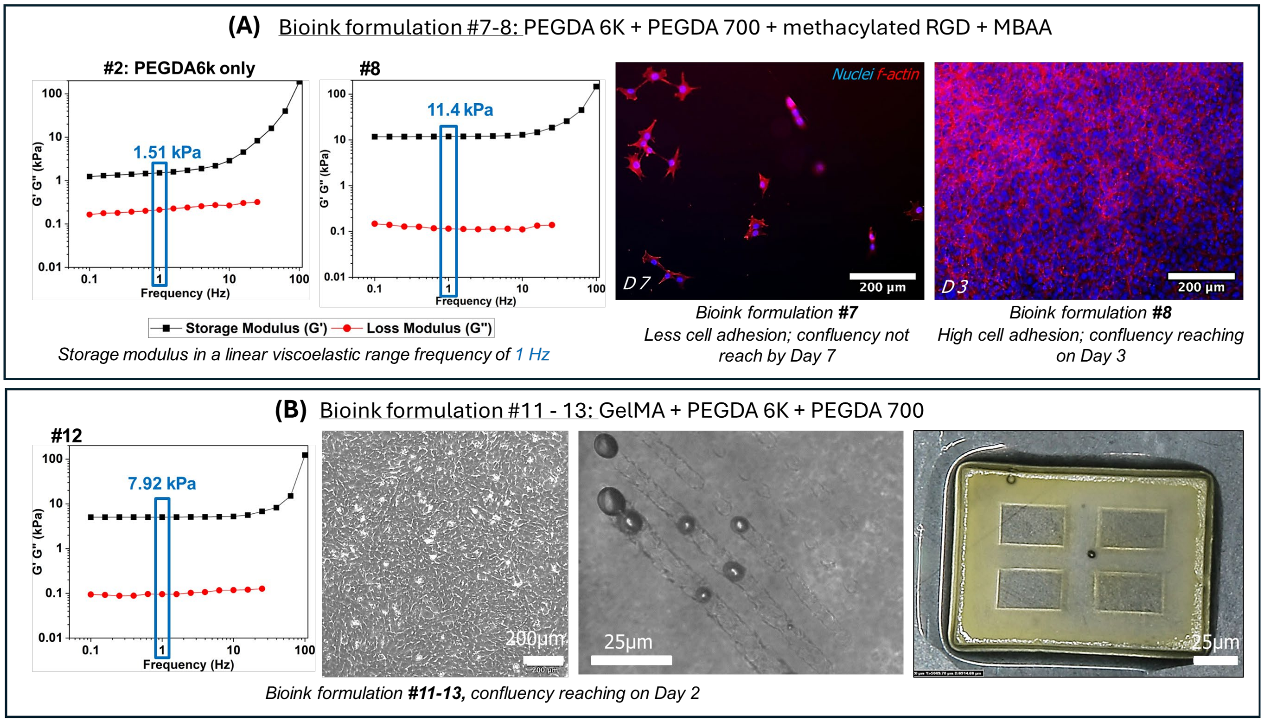


**SI-3.** (A). Rheological analysis of bioink formulations #7, #8. The storage modulus (G') was measured within the linear viscoelastic range at a frequency of 1 Hz. Fluorescence images showing cell morphology. Cells reach 100% confluence on Day 3 for formulation #8 as compared to formulation #7. (B). Rheological analysis of bioink formulation #12, showing cell confluence on Day 2. Representative images show fidelity of TPA-mode and DLP-mode using formulation #12. Qualitative screening was performed using 2H11 cell line.

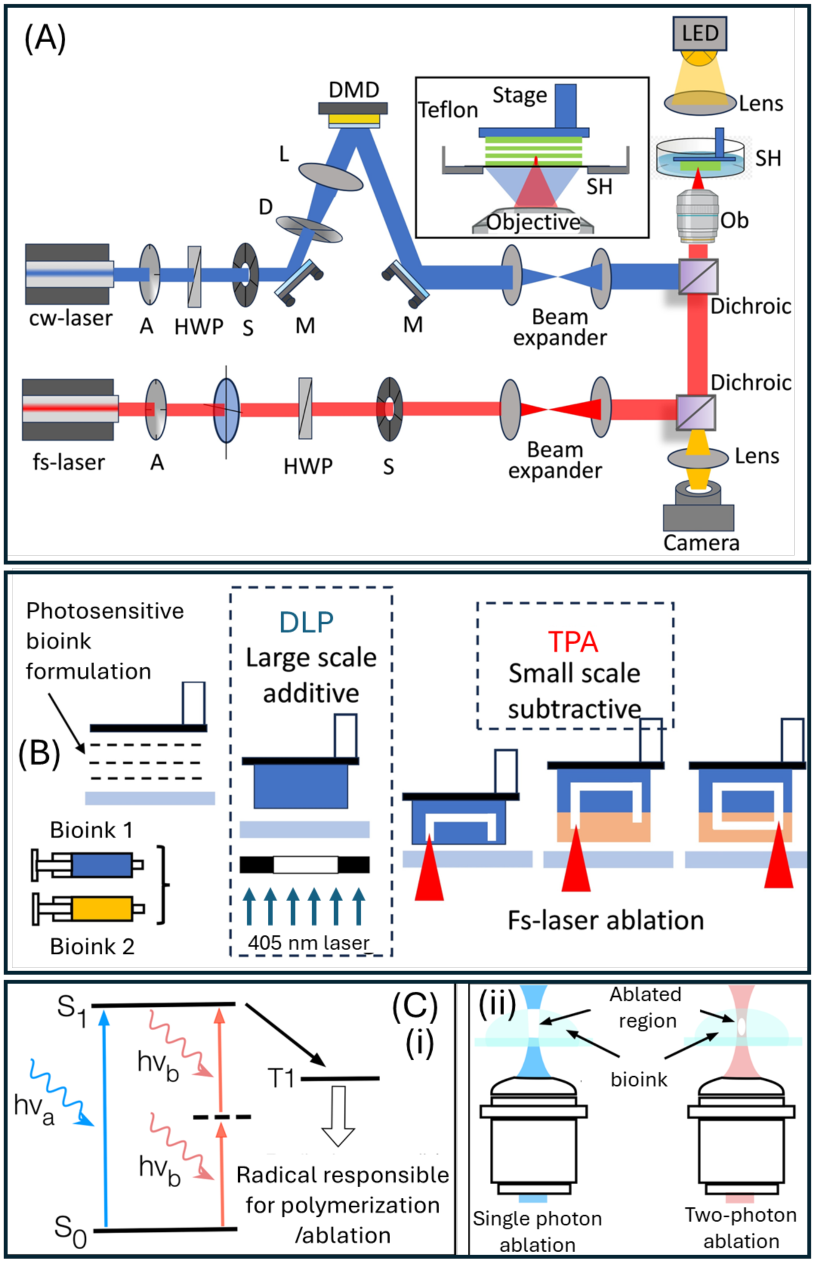


**SI-4.** (A) Schematic of a hybrid additive-subtractive platform. [fs- laser—femtosecond laser, cw-laser-continuous wave laser, A –attenuator, HWP—half waveplate, S- shutter, Ob- Objective lens , SH-sample holder, M- Mirror, D- Diffuser, L-Lens, DMD- Digital micro mirror device, LED-Light emitting diode, DLP-Digital Light processing, TPA- two photon ablation]. (B) Schematic of process flow of sequential and bidirectional additive DLP-mode and subtractive TPA-mode. (C) (i)Jablonski diagram showing the two-photon absorption processes and depiction of the formation of radical responsible for polymerization formed in the triplet state. (ii) Multiphoton ablation relies on the simultaneous absorption of two-photon near-infrared light, enabling high-resolution 3D control while single-photon ablation is mostly restricted to planar fabrication.

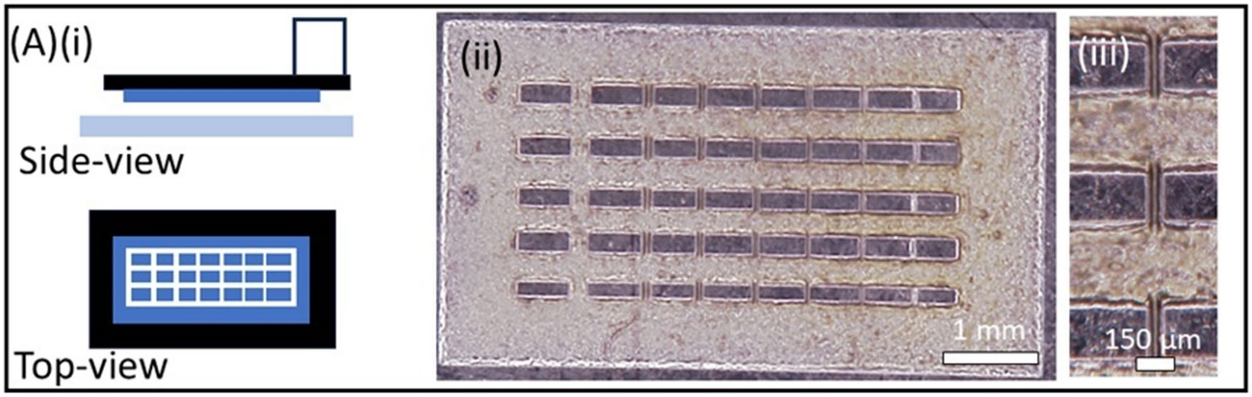


**SI-5.** (A) Schematics of side and top views of the structure used to test lateral (XY) resolution of DLP-mode (ii) Printed structure and (ii) a zoomed-in view. Laser power = 280 mW, Exposure time = 2 seconds.

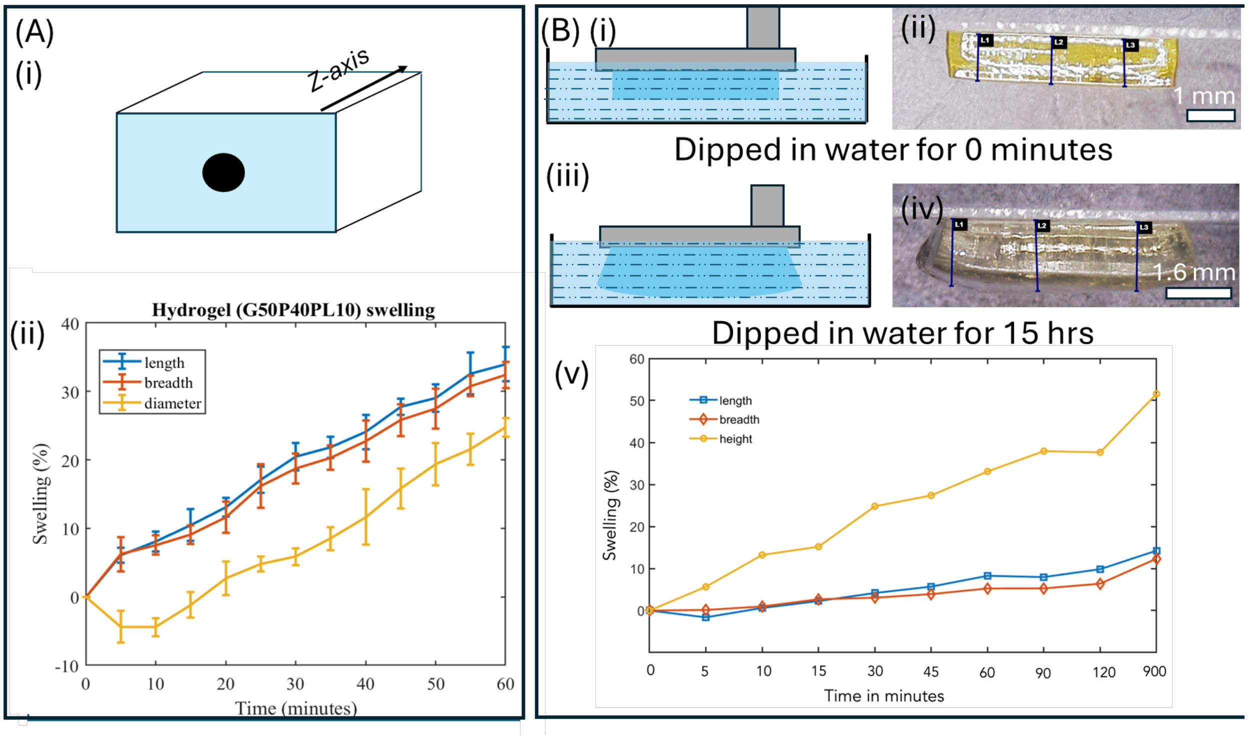


**SI-6.** (A) A rectangular slab featuring a vertical hole was fabricated using DLP-mode (i) Schematic of the slab. (ii) Dimensions of printed slabs, detached from the coverslips, were measured using a HIROX microscopy, followed by immersed in a water bath and allowed to swell at room temperature. Plot showing the changes in the slab dimensions over a period of 60 minutes. (B) Since during TPA-mode, the crosslinked structure remains attached to the coverslips, we investigated the swelling behavior under constrained condition (i-iv) Schematic representation and DLP-printed slab attached to an L-shaped stage, undergoing swelling. (v) Plot showing the changes in the slab dimensions: **Note:** All work was conducted using bioink formulation #12. Results show an increase of 35% in the z-direction and an increase of 10% in the lateral (XY) direction. The mismatch in swelling ratios between Z and XY directions, due to constraints provided by the methacrylate coverslips, results in the formation of a slightly dome-shaped top surface (**iv**).

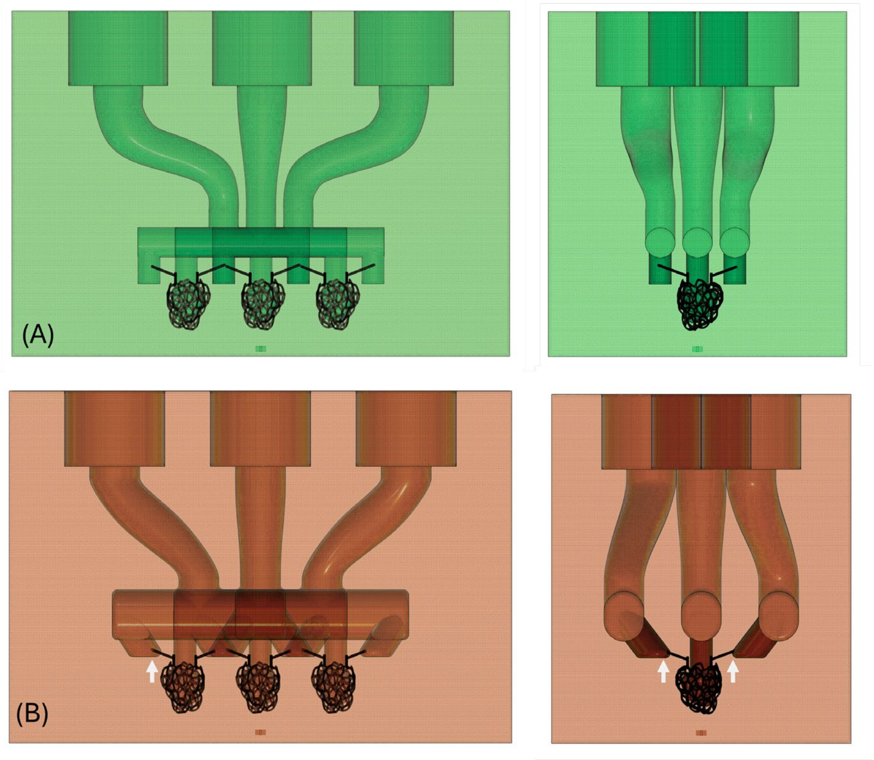


**SI-7. Modification of CAD design to facilitate perfusion of media into 3D capillary-like network structure.** (A) Abrupt transition from meso-scale channels to microchannels results in unreproducible perfusion. (B) Gradual transition of cross-sectional, as shown by white arrows, result in reproducible perfusion flow in structurally complex capillary networks.

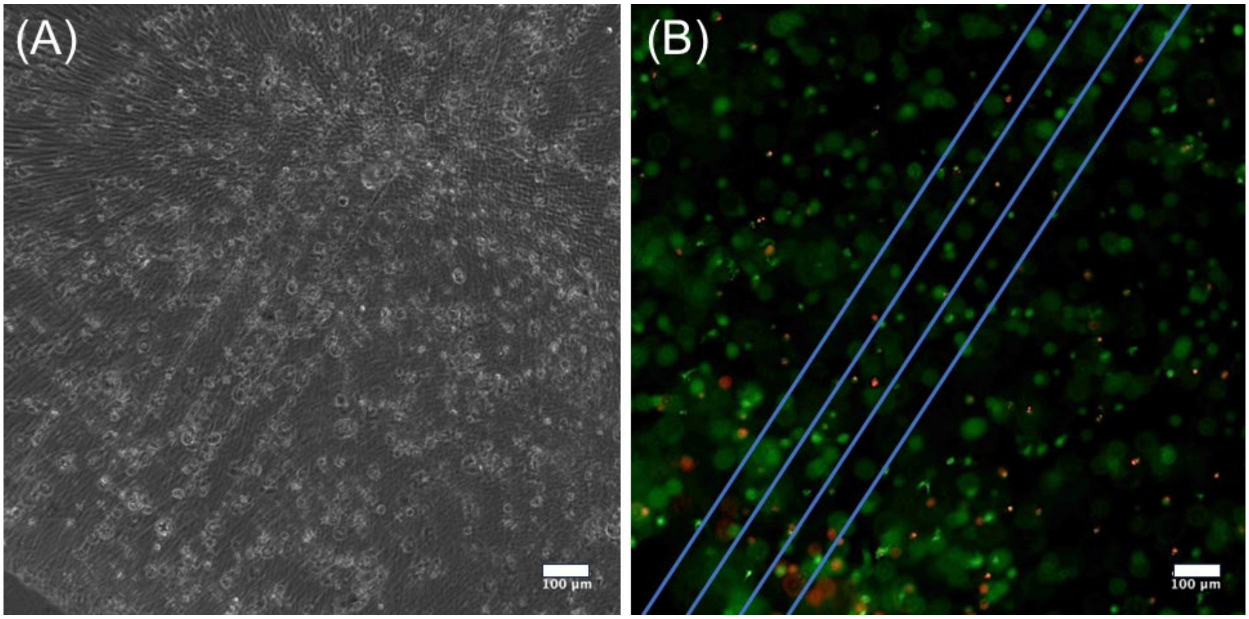


**SI-8.** DLP-mode was used to print 10T1/2-laden bioink slab using formulation #12 followed by use of TPA-mode to ablate an array of microchannels of size ~20µm. Representative brightfield and fluorescence images are shown in A and B.

**Video Captions**

V1. Video showing femtosecond laser–induced cavitation in in the crosslinked slab structure generated using DLP-mode and bioink formulation #12. TPA-mode generates bubbles which expand during laser irradiation and collapse once the laser beam is switched off.

V2. Video showing that during laser scanning, the cavitation bubble follows the laser focal point, creating a hollow microchannel.

V3. Video showing perfusability of microchannels (lumen size of 10µm) embedded ~250µm inside bioink chip using a red microbead solution (1.0 µm FluoSpheres™, Invitrogen).
